# Highly Robust DNA Data Storage Based on Controllable GC Content and homopolymer of 64-Element Coded Tables

**DOI:** 10.1101/2023.09.27.559852

**Authors:** Lu Yunfei, Zhang Xuncai

## Abstract

In this paper, we propose a DNA storage encoding scheme based on a 64-element coding table combined with forward error correction. The method encodes the data into DNA sequences by LZW compression of the original text, adding error correction codes and scrambling codes. In the encoding process, the effects of GC content limitation and long homopolymers on DNA sequences are considered. At the same time, RS error correction code is introduced to correct the DNA sequence to improve the accuracy of decoding. Finally, the feasibility and effectiveness of the program were verified by simulation experiments on Shakespeare’s sonnets. The data results show that the GC content of DNA sequences encoded by the program is kept at 50%, the homologous multimer length is not more than 2, and the original information can be recovered from the data of 10-fold sequencing depth without error with an error rate of 0.3%. We conducted simulation experiments of primer design, DNA sequence recombination, PCR amplification, and sequence reading on DNA sequences loaded with design information, which further proved the concrete feasibility of the scheme. This scheme provides a reliable and efficient encoding scheme for DNA information storage.

## 1. Introduction

Traditional physical storage media can no longer meet the exponentially growing demand for data storage. According to forecasts, the global data volume will increase to 1.75×10^14^ GB by 2025[1, 2]. Data storage problems have been solved using information storage in organic molecules such as DNA molecules, oligopeptides, and metabolite moieties[3]. Compared with traditional storage media, these emerging storage media show significant advantages in terms of storage density, especially the unique double helix structure of the DNA molecule. The storage capacity can be as high as 455 EB bytes per gram of DNA molecules. In addition, DNA molecules have a half-life of about 521 years under proper storage conditions, Half-life above 2 million years if stored in silica[4]. Based on the advantages of DNA molecules, such as high storage density and long-term stability, this ancient information carrier is regarded as a storage medium with great potential. In the DNA data storage process, the raw information is first converted into a binary sequence and then into a DNA sequence according to specific coding rules. These DNA sequences can be synthesized and stored in organisms or in vitro as oligonucleotides or double-stranded DNA, facilitating subsequent retrieval and reading of the original information using relevant technologies. Thanks to its rapidity and accuracy, the latest third-generation DNA molecular sequencing technology makes it easier to read and write the information content stored in DNA molecules[5]. In conjunction with third-generation DNA molecular sequencing technologies, several random read strategies have been proposed to achieve selective access to stored information[6-8], further enhancing the utility and scalability of DNA data storage.

DNA sequences follow biosynthetic and sequencing constraints, effectively reducing errors during subsequent reads and increasing decoding efficiency[9]. Homopolymer (nt) and GC content (% calculation) are two critical metrics for assessing the performance of coding schemes. For example, single base long string repeats (homopolymers) of lengths greater than five may introduce higher error rates during synthesis or sequencing[10, 11]. In addition, GC contents below 40% or above 60% (i.e., extreme GC contents) are usually detrimental to the synthesis of DNA molecules[11-14]. Church[15], Grass[16], Blawat[17], and Erlich[18] used a binary transcoding strategy (0 for A or G, 1 for C or T) to control the GC content to avoid long homopolymer sequences. Subsequently, it has been shown in the literature[19] that LT codes can efficiently handle many input symbol segments (also known as droplets), effectively transmitting information in erasure channels. DNA Fountain uses LT codes as an internal transcoding strategy, which significantly reduces the introduced redundancy. DNA Fountain screens droplets that meet the constraints according to their unique screening mechanism to avoid extreme GC content and long homopolymers. However, the successful decoding of LT codes relies on introducing sufficient logical redundancy, which will lead to decoding failure if too few droplets meet the criteria[18, 19]. Reducing logical redundancy may increase the decoding failure probability while increasing analytical tedium decreases the information density and the synthesis cost.

At the same time, the transcoding method using LT code as the internal code is also rigorous in selecting the input symbol segment length. If the input symbol segment length and the message eBCH code length are chosen arbitrarily, it will lead to rate degradation in the fading channel[20]. To ensure the integrity of the information, Goldman used a quadruple physical redundancy approach, where an excess number of DNA molecules are synthesized, increasing the number of copies of DNA molecules per sequence[21]. However, this can increase the cost of subsequently synthesizing the DNA sequences. Therefore, when developing transcoding algorithms, they need to be characterized by high storage density and high robustness to adapt to various types of data transcoding[22], and at the same time, the cost should be reasonably controlled to apply the DNA information storage in practical scenarios.

To this end, this paper proposes a DNA storage transcoding scheme based on a 64-element coded mapping table. First, the text to be read and written is compressed, and an RS error correction code is added to ensure data integrity[18, 23-25]. Then, a perturbation sequence is introduced to prevent consecutive occurrences of the same binary element. Finally, the data is transcoded into DNA base sequences according to the proposed mapping table.

## 2. Restrictions on DNA coding

### 2.1 Hamming distance constraints

The Hamming distance is a measure of similarity between two encoded code words. In DNA coding, a smaller Hamming distance means that two DNA strands have more of the same number of bases between them. For two DNA sequences, a and b, we use *H(a, b)* to denote the number of different bases at position *i* of the sequence *(a, b)*. The Hamming distance is calculated using the following formula[26]:

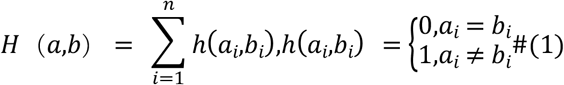

Where *H(a, b)* denotes the Hamming distance between two bases, *0* when the corresponding bases are the same, and *1* when the complementary bases are different.

### 2.2 GC content constraints

GC content refers to the ratio of the total number of G and C bases to the total number of bases in a DNA sequence, which is closely related to the stability of the DNA sequence and the melting point. The designed DNA sequence should be within the range of *40% ≤ GC(n) ≤ 60%* to ensure that the constraints of GC content are met, and the formula of *GC(n)* is as follows[26]:

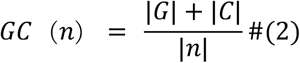

where *GC(n)* denotes the GC content of the sequence, |*G*| and |*C*| represent the number of bases G and C, respectively, in the sequence *n*, and |*n*| denotes the total number of bases in the sequence.

### 2.3 Homopolymer constraints

DNA sequences should satisfy the occurrence of longer repetitive base sequences (homopolymers) that do not contain them. Longer consecutively repeated bases can lead to misinterpretation of the synthesized DNA sequence during subsequent sequencing, thus reducing the robustness of the decoding. For example, in the sequence *TCCCCAC*, the C bases are repetitive, making it easy to read long *C* as short *C* during synthesis and sequencing, increasing the error rate in the DNA storage information and decreasing the accuracy of reads and writes. This can be avoided by limiting the length of the homopolymer. For example, *AAA* is a shorter homopolymer that is controlled to be between 3 and 5 bases long, which can be read accurately by third-generation sequencing instruments.

For the codeword *d(D*_*1*_, *D*_*2*_, *D*_*3*_, …, *D*□), where *D*□ *∈ (A, T, G, C)*, for any *i*, we have the following relation[26]:

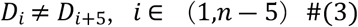

## 3. DNA encoded storage program design

An ideal transcoding scheme should satisfy the constraints of Section 2 and be able to correct insertion and deletion errors. To this end, we propose a transcoding method based on a 64-element coded table. In this scheme, the original text is first compressed using the LZW mechanism and converted into a binary stream. An RS check digit block is added to the binary stream to generate a binary stream with an error-correcting code. The binary stream with an error-correcting code is scrambled, and finally, the rearranged binary sequences are transcoded into DNA sequences using a 64-element coded table. The steps are shown in Figure 1:

**Fig. 1.**
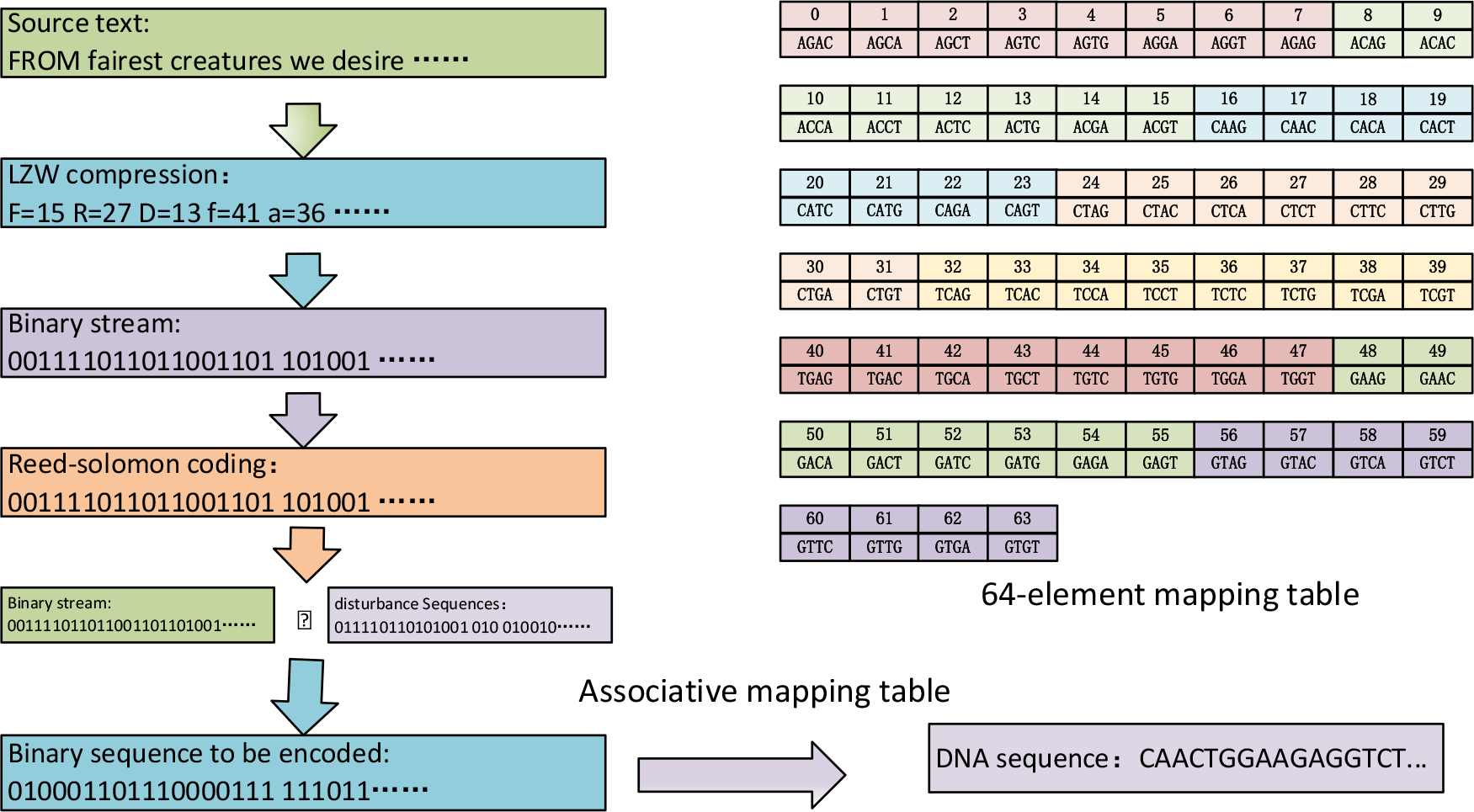
Flowchart of coding process of 64 elements coded table

### 3.1 Design of the 64-element coded table

The DNA information storage process consists of four main steps[27]: (a) Encoding digital information into a DNA sequence. (b) Synthesizing DNA sequences. (c) Storing the DNA sequence. (d) Recovering the stored DNA information by sequencing. However, current high-throughput sequencing technologies (e.g., Illumina sequencing methods) may lead to complex pairing of bases during amplification of DNA sequences, which may trigger base mutations, produce long homopolymers that interfere with the information carried by the original DNA sequence, and cause misinterpretation in subsequent sequencing processes[28]. Therefore, an excellent transcoding scheme should satisfy the following constraints during DNA information storage: 1) avoiding long homopolymers and 2) avoiding extreme GC content. The scheme selects 64 combinations from 4^4^=256 base pairs that meet the GC content and homopolymer constraints to form a 64-element coded table.

In the process of converting binary code streams to DNA sequences, we note that overly long binary code streams mapped to shorter DNA base sequences are irreversible, and excessively short binary code streams mapped to longer DNA base sequences result in a decrease in the information density and an increase in the cost of synthesis. Thus, we consider every four base alignments in the set of 4^4^ = 256 codes as a set of mapping elements and filter these mapping elements by considering the limitations of homopolymers. For example, if the previous set of mapping elements is *AAAA*, the next set of mapping elements cannot have *AAAA* again to avoid longer homopolymer sequences. Also, while considering longer homopolymer sequences, we note that extreme GC content occurs when transcoding into DNA sequences if there are too many occurrences of *G* and *C* bases in each set of mapped elements. For example, a set of coding features with a *GCGC* has a GC content of up to 100%, which can lead to breaking hydrogen bonds and denaturation difficulties during the synthesis of DNA sequences. Therefore, we filtered the mapping elements with GC content of less than 25% or more than 75% in the coding set and excluded them from the coding set. After screening, we obtained 64 code mapping tables that meet the requirements, as shown in Table 1 below:

**Table 1.** 64-element coded mapping table.

|  |  |  |  |  |  |  |  |  |  |  |  |  |  |  |  |
| --- | --- | --- | --- | --- | --- | --- | --- | --- | --- | --- | --- | --- | --- | --- | --- |
| 0 | 1 | 2 | 3 | 4 | 5 | 6 | 7 | 8 | 9 | 10 | 11 | 12 | 13 | 14 | 15 |
| AGAC | AGCA | AGCT | AGTC | AGTG | AGGA | AGGT | AGAG | ACAG | ACAC | ACCA | ACCT | ACTC | ACTG | ACGA | ACGT |
| 16 | 17 | 18 | 19 | 20 | 21 | 22 | 23 | 24 | 25 | 26 | 27 | 28 | 29 | 30 | 31 |
| CAAG | CAAC | CACA | CACT | CATC | CATG | CAGA | CAGT | CTAG | CTAC | CTCA | CTCT | CTTC | CTTG | CTGA | CTGT |
| 32 | 33 | 34 | 35 | 36 | 37 | 38 | 39 | 40 | 41 | 42 | 43 | 44 | 45 | 46 | 47 |
| TCAG | ACAC | TCCA | TCCT | TCTC | TCTG | TCGA | TCGT | TGAG | TGAC | TGCA | TGCT | TGTC | TGTG | TGGA | TGGT |
| 48 | 49 | 50 | 51 | 52 | 53 | 54 | 55 | 56 | 57 | 58 | 59 | 60 | 61 | 62 | 63 |
| GAAG | GAAC | GACA | GACT | GATC | GATG | GAGA | GAGT | GTAG | GTAC | GTCA | GTCT | GTTC | GTTG | GTGA | GTGT |

### 3.2 Adding a Perturbation Sequence

Information scrambling is a standard method used in communications engineering to ensure the security of signal transmission and copyright protection of transmitted information against illegal interception and theft. In this paper, the identification of the above parameters of demodulation of the position error signal is achieved by two improvements: adding the voltage injection vector and optimizing the signal demodulation. Adding artificial noise processing is a standard scrambling method in satellite signal transmission[29]. There are two methods of signal scrambling: one is to run the coded signal sequence by superimposing pseudo-random lines, and the other is to use cryptographic algorithms (e.g., DES) to encode the digitized coded signal for transmission in segments. During DNA storage, Literature[26] and literature[30] state that, Introducing a pseudo-random sequence to randomize the input data can effectively break the case of multi-bit repetitions (e.g., 011111111110).

### 3.3 Single DNA sequence design and error correction

The complete DNA sequence includes not only information loading regions and error correction regions but also indexing regions in the DNA sequence to realize the need for fast access to information[6]. According to the previously described processing for Shakespeare’s sonnets, the original text, totalling 97,343 bytes, was compressed, error corrected, and scrambled, resulting in a 399,408-bit binary code stream, ultimately synthesizing 1,841 DNA sequences. To ensure complete retrieval of each DNA sequence, we designed the length of the information load region and the RS error correction region to be 129 base pairs plus 15 base pairs, totalling 144 base pairs (217 bits). Each DNA sequence is designed with a 16-base index region to enable random access or direct readout. After testing, data less than 0.47 GB can be quickly located and accessed through the index area. In addition, primer regions were added to the DNA strand to facilitate data backup and sequencing builds. As the Illumina sequencing method required, primers of 20 bases were added to each end of the single-stranded DNA sequence for library amplification.

During the encoding process, we incorporate error-correcting regions to improve the stability of the data to prevent errors caused by base mutations. The dominant error-correcting codes include Hamming error-correcting codes[31] and RS error-correcting codes[16, 32]. In contrast, Hamming error correction codes can only correct one error or detect multiple errors, while RS error correction codes can convert any number of mistakes[30, 31]. In choosing the encoding scheme, we wanted to ensure the integrity of the digital information while reducing the total number of DNA bases to minimize the cost of synthesis. Considering the fidelity of the DNA sequence and the synthesis cost issues, the RS error correction code was chosen as the error correction method in the encoding process in this scheme. Each single-stranded DNA sequence fragment is designed as follows:

According to a 2016 study[17], substitution error rates accounted for a more significant percentage of errors, with insertions and deletions being less likely. Substitutions cause most errors, so the theoretical coding scheme should be able to handle substitution errors. During experiments, extreme GC content and longer homopolymer stretches (e.g., long homopolymer AAAAA) can lead to substitution errors during synthesis and sequencing. Literature[17] and literature[26] point out that the substitution error rate increases significantly, especially for homopolymer sequences of lengths more than six. In addition, DNA sequences with high GC content develop a highly variable state during PCR amplification. Literature[33] also indicated that the coefficient of variation reached 11.8% in five separate sequences in a population of 20 mimics sequenced for the Ion Torrent PGM 16S rRNA gene. This increase in the coefficient of variation is due to differences in the GC content of the base sequences leading to increased variation during PCR. Once this difference occurs, the number of insertion, substitution and deletion errors in the base sequences increases. In the face of mutations triggered by extreme GC content and substitution errors started by long homopolymers, the 64-element coded table proposed in this paper has potential advantages for error correction. Simple substitution, deletion, and insertion errors can be corrected according to this coding table, as shown in Fig. 3. The three error cases are briefly described below:

**Fig. 2.**
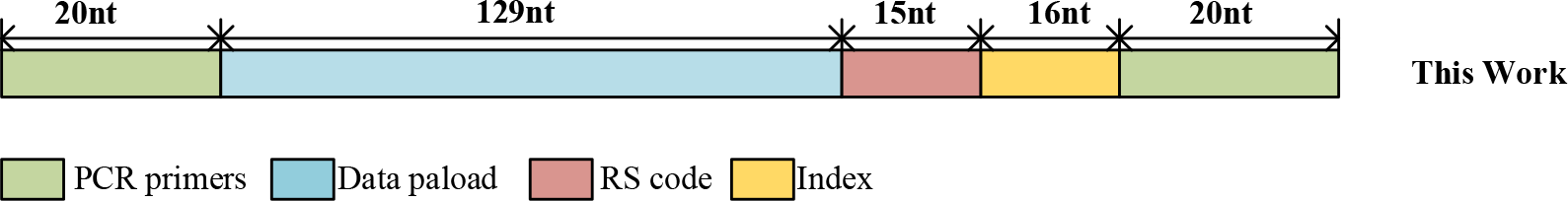
Sequence design of oligonucleotides designed in this paper

**Fig. 3.**
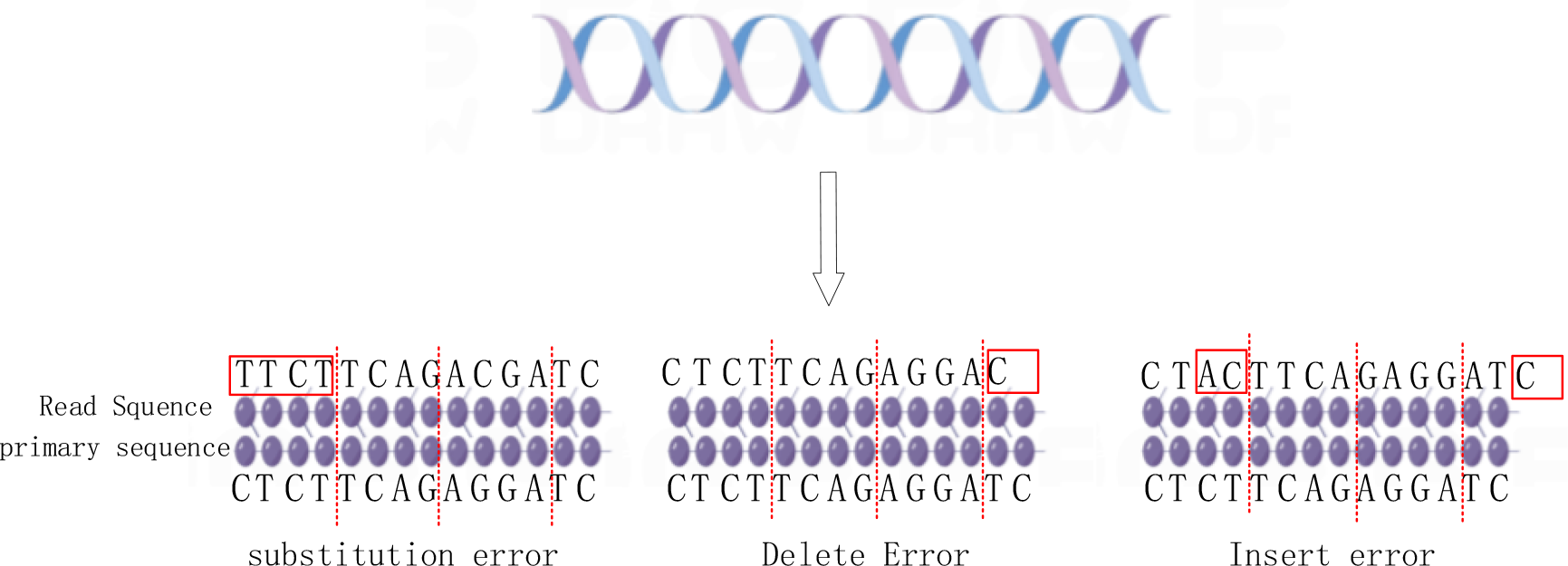
Self-correcting replacement, deletion and insertion errors

#### Case 1: Replacement error

According to the above encoding, read the DNA sequence such as *CTCTTCAGAGGATC* …… In the traversal process, every 4 bp as a group, if there are *TTCT, CCAG* such as not in the 64-element coded table mapping elements, you can judge that a substitution error has occurred, and according to the 64-element coded table to find the correct mapping elements to be corrected.

#### Case 2: Deletion error

The DNA sequence was read based on the above encoding, such as *CTCTTCAGAGGATC* …… In the traversal process, every 4 bp as a group, the first few groups of DNA sequences did not occur error, but in the last group of base sequences, only *C* base appeared. After comparison, the DNA sequence length is shorter than the designed DNA sequence, so it can be judged that the deletion error occurred.

#### Case 3: Insertion error

The DNA sequence is read based on the above encoding, such as *CTCTTCAGAGGATC* …… During traversal, the DNA sequence is divided into groups every 4 bp. In the read results, if the second pair of bases in the previous set is *CT*, and in reading the first pair of bottoms in the subsequent group, the original DNA sequence is *CT*. Still, the read sequence appears as *AC*, indicating that an error has occurred. Then continue traversing the subsequent bases until the end. By combining this with the 64-element coded table, we can find that an error has occurred before the AC mapping element and continue traversing until the end. When performing sequence comparison, if the number of bases in the read DNA sequence is more than the number of floors in the designed DNA sequence, it can be determined that an insertion error has occurred. However, based on the 64-element coded table, only insertion errors can be detected, and the location of the error is determined, but no correction can be made based on the site. This situation is similar to a deletion error, and it is impossible to accurately determine the actual problem of the group of base sequences to determine the specific correction of the insertion error.

The above is a brief description of the three error scenarios. The static coding table of this scheme has some self-correcting advantages for substitution-type errors. However, for insertion and deletion type errors, this coding table can only detect the error but not correct it. We introduce a simpler error correction code to avoid the situation where insertion and deletion errors lead to decoding failure. Specifically, we update each DNA base sequence using the RS(255, 240) error correction code to ensure the fidelity of the DNA sequence[16].

By introducing RS (255, 240) error-correcting codes, we were able to correct enough errors to improve the reliability of the DNA sequence. RS error-correcting code is an error-correcting code scheme capable of repairing any number of mistakes. In our project, applying RS (255, 240) to each DNA base sequence corrects a certain number of insertion and deletion errors, which enhances the accuracy and stability of decoding.

By combining static coding tables and RS error-correcting codes, we can handle both substitution-type errors and correct a certain number of insertion and deletion errors, thus improving the overall decoding capability and data fidelity.

## 4. Results and analysis

In the previous literature, some computational methods and corresponding concepts related to information density have been proposed for evaluating the performance of coding schemes. Among them, base coding density (bit/nt) describes the number of effective bits carried by a base in practice, regardless of the effect of molecular copy number[15, 18, 34].

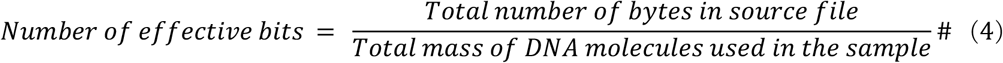

In addition, several concepts have been proposed, such as coding potential and realized capacity, which involves complex theoretical derivations in information theory. However, in DNA storage, these concepts have not been recognized on a large scale due to the wide variation of data in practical applications.

Our study used a DNA sequence length of 129 bases per segment as the information carrier, corresponding to 194 bits of information, and calculated storage density. We have improved the net coding efficiency by using the LZW mechanism compression on the original text. Eventually, we obtained a net information density of 1.49 bit/nt.

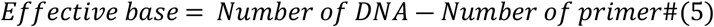

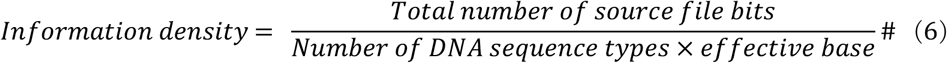

This program takes Shakespeare’s sonnets as an example for a reading test, and the specific steps are as follows:

a. Shakespeare’s sonnets (97,343 bytes in the original text) were first LZW compressed, resulting in 24,963 compressed text information.
b. To ensure the text message’s recovery and the original message’s fidelity, we converted 24963 bytes to a binary stream according to ASCII code and split every 240 bits into segments. Then, 15 redundant bits are added to each fragment to correct the error of 10 bits.
c. After error correction coding, the total length of the binary code stream is 399408 bits.
d. The binary code stream is scrambled with chaotic sequences. After the scrambled sequence is added to the binary data stream, every 3 bytes are encoded into 16 bases (8 bits per byte for 24 bits) according to the 64-element mapping table.

Ultimately, Shakespeare’s sonnet was transformed into 266,272 bases. Based on the above calculations, this scheme achieves a theoretical storage density of 399408/266,272 = 1.5 bits/base. Combined with the GC content involved, we summarize and list Table 2 below:

**Table 2.**
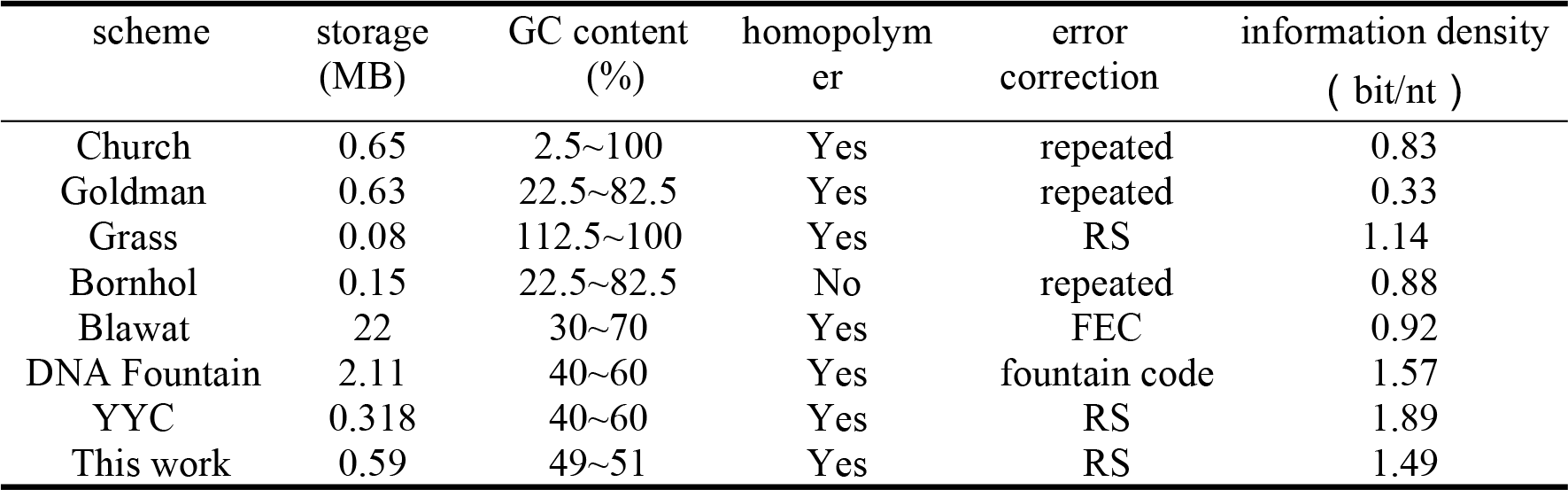
Comparison with previous work.

| scheme | storage<br>(MB) | GC content<br>(%) | homopolym<br>er | error<br>correction | information density<br>( bit/nt ) |
| --- | --- | --- | --- | --- | --- |
| Church | 0.65 | 2.5~100 | Yes | repeated | 0.83 |
| Goldman | 0.63 | 22.5~82.5 | Yes | repeated | 0.33 |
| Grass | 0.08 | 112.5~100 | Yes | RS | 1.14 |
| Bornhol | 0.15 | 22.5~82.5 | No | repeated | 0.88 |
| Blawat | 22 | 30~70 | Yes | FEC | 0.92 |
| DNA Fountain | 2.11 | 40~60 | Yes | fountain code | 1.57 |
| YYC | 0.318 | 40~60 | Yes | RS | 1.89 |
| This work | 0.59 | 49~51 | Yes | RS | 1.49 |

### 4.1 Single DNA sequence

Scrambling during data transmission aims to randomize the original binary code stream. To achieve this goal, in the binary code stream, after adding the error correction code, we employ a specific chaotic sequence for scrambling and descrambling it before proceeding to decode. Precisely, we introduce logically mapped chaotic sequences with expressions such as Eq. (7):

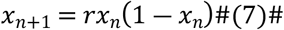

Among them, for the case of *3*.*569945627 < r ≤ 4*, we take advantage of the characteristics of logical chaotic sequences, such as high sensitivity and unpredictability, and use the arrangements as mentioned above and the binary code streams after adding the error correction code for the dissimilarity processing, to avoid the occurrence of too many consecutive {0, 1} in the binary code streams, and thus prevent the occurrence of more extended consecutive identical bases in the subsequent synthesis of DNA base sequences in the following DNA base sequence synthesis. In addition, the scrambling process can also play a role in encryption.

Figure 4 illustrates the percentage of {0, 1} elements before and after the sequence scrambling and the degree of chaos of the series at different cycles. According to the observation in Fig. 4(a), the distribution density of *X(n)* values produced by the logistic mapping sequence between *3*.*7 < r ≤ 4* is much higher than in the case of *2*.*5 < r ≤ 3*.*7*. Therefore, we chose the chaotic sequence generated by *r = 3*.*8* and the given parameter *X*_*0*_ *= 4*.*0* for heterodyne processing with the binary code stream. After scrambling, the number of 0s and 1s in the binary sequence is relatively uniformly distributed, around 50%.

**Fig. 4.**
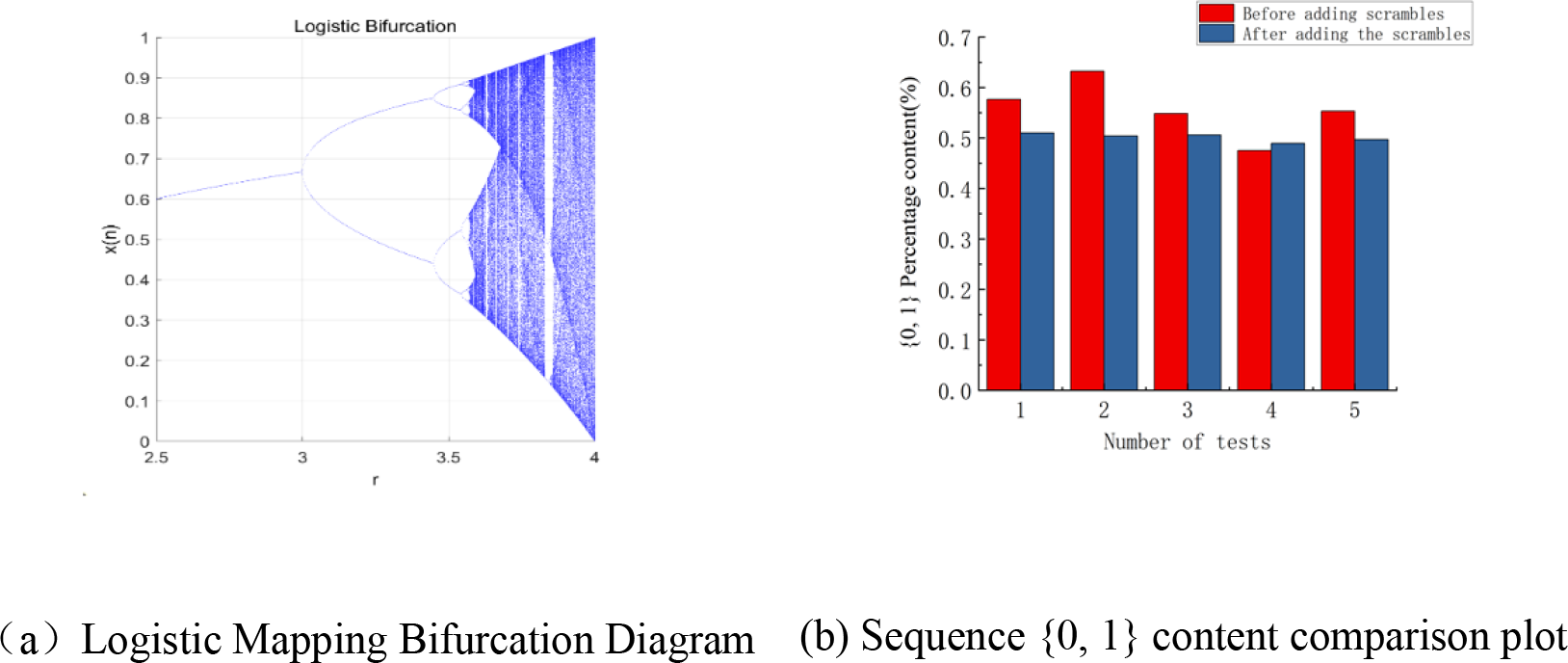
Bifurcation map of logic mapping and comparison of {0, 1}

### 4.2 Code snippet performance analysis

#### 4.2.1 GC content analysis

Higher GC levels can make it challenging to break hydrogen bonds, making DNA denaturation difficult. To raise the melting point to break the hydrogen bonds, an increase in temperature increases the activity of the DNA bases, which increases the probability of mutation of the DNA bases, leading to a rise in the error rate during DNA sequencing. Therefore, these factors need to be weighed when controlling the GC content.

We tested this scheme and several other transcoding methods on five documents and compared their GC content. We used a sequencing depth of 10× for each text and calculated the average GC content percentage for each transcoding scheme. The results are shown in Fig. 5(a)∼(e).

**Fig 5.**
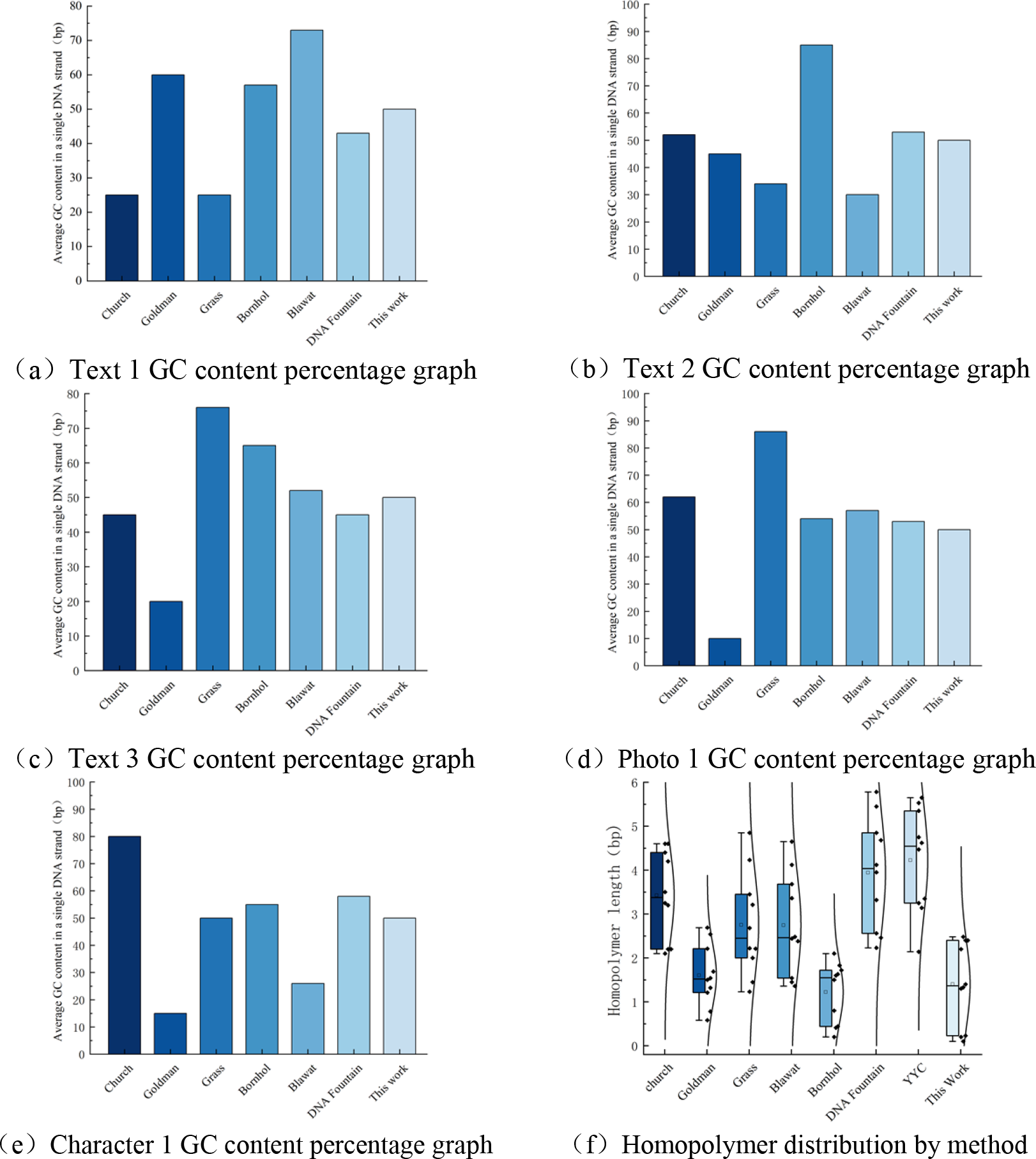
Percentage of GC content and homopolymer distribution by method

The transcoding scheme of DNA Fountain[9] controls the average percentage of GC content between 40% and 60%. The transcoding method of Church[15] keeps the average rate of GC content between 25% and 80%. The transcoding scheme of Grass[16] slightly decreases the average percentage of GC content compared to Church’s transcoding strategy, controlling it between 25% and 78%, but did not fully comply with the requirements of the sequencing method. The transcoding scheme of Goldman[35] controlled the average percentage of GC content between 15% and 60%, which is still a gap compared with the DNA Fountain and YYC codes.

Considering the requirements of the third-generation sequencing system for GC content and the constraints of upstream and downstream biological experiments, this program controls the average percentage of GC content between 49% and 51%, which better adapts to the requirements of the third-generation sequencing method in biological experiments, this allows a small number of substitution, deletion and insertion errors of DNA bases during sequencing.

Considering the requirements of the third-generation sequencing system for GC content and the constraints of upstream and downstream biological experiments, this program controls the average percentage of GC content between 49% and 51%, which better adapts to the requirements of the third-generation sequencing method in biological experiments, this allows a small number of substitution, deletion and insertion errors of DNA bases during sequencing.

#### 4.2.2 Homopolymer analysis

The most accurate third-generation sequencing technologies currently available cannot read DNA homopolymer sequences entirely and correctly and are especially difficult to discriminate between multiple consecutively repeated bases. In the methods of this paper, we tested the stored document Photo 1 (Cameraman 256) at 10× sequencing depth by minimizing the homopolymer length. We compare the performance of several different transcoding schemes in terms of homopolymer length distribution (denoted as length n). The specific results are shown in Fig. 5(f).

For the transcoding schemes of DNA Fountain[9] and YYC[23], most of the homopolymer lengths are clustered at n=4, which is long compared to other transcoding methods. This is because they belong to dynamic coding, which cannot control the generation of the last base, which is a disadvantage of dynamic coding. As for Blawat[17] and Bornholt[36] transcoding methods, most homopolymer lengths are concentrated at n=1∼2. In Grass[16] transcoding scheme, most homopolymer lengths are at n=2∼3, and the most extended homopolymer length is at n=4.

After comparison, our method successfully controls the homopolymer in the range of homopolymer lengths of n=1∼2, which can satisfy the homopolymer constraints in the sequencing process.

### 4.3 Read Process Performance Analysis

In Shakespeare’s sonnet simulation experiments, we constructed coding tables by randomly selecting 64 mapping elements from a set of 256 codes. We then compared the base substitution error rates between (*G, C*) and (*T, G*) using the 64-element coded table with those of the random coding table, as shown in Fig. 6(a), (b). The results show a 1.08% decrease in the substitution error rate between (*G, C*) and an 18.01% decrease in the substitution error rate between (*T, G*) when using the 64-element coded table. However, the base substitution error rate between (*T, C*) and (*A, C*) increased slightly, but not more than 10%. The details are shown in Fig. 6(c). Using the 64-element coded scheme proposed in this paper has a lower base substitution error probability than the random coding table.

**Fig. 6.**
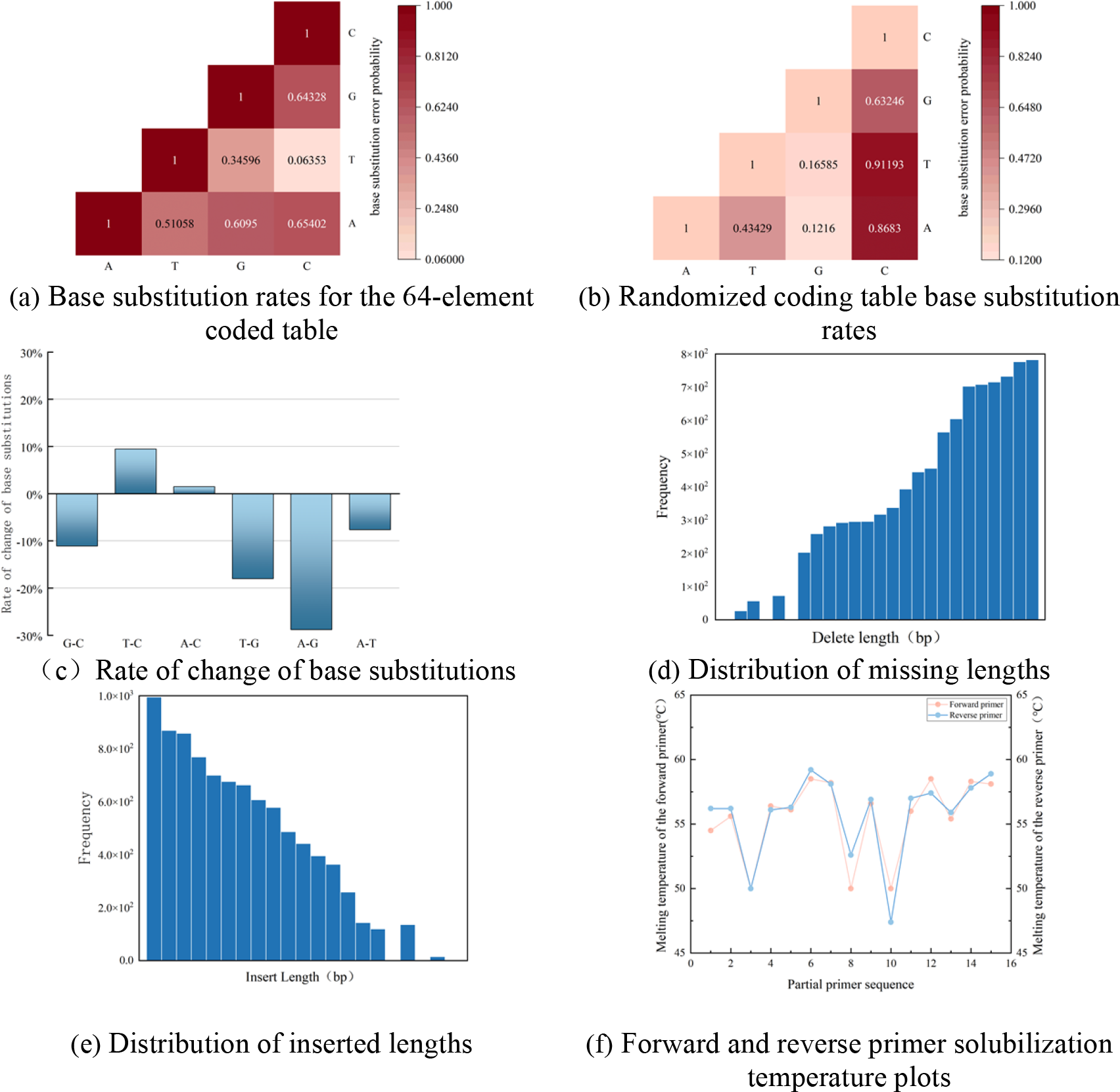
Various error analyses and forward and reverse primer solubilization temperature plots

We used the Flye[37] platform to assemble from scratch the DNA sequences generated by the complete Shakespearean sonnets. These sequences generated 266,272 bases according to a 64-tuple coding table (note that we only analyzed errors in the informational regions and did not encode error correction in such areas as the index). We split these DNA sequences into informative loads and RS error-correcting fragments (129 nt + 15 nt), yielding 1,850 DNA strands, which were then recovered. We added pseudo-random errors in the generated DNA sequences (equal number of insertions and deletions). We found that the error case of losing 200 bases occurred less than 1% of the time, and the error frequency of inserting 20 commands was almost close to 0. Thus, the coding method based on the 64-element coded mapping table ensures highly robust recovery.

### 4.4 Monolithic Segment Read Performance Analysis

When recovering stored data, retrieving the information carried by the DNA sequences in the information payload region is crucial. As a maximum distance separable code, the Reed-Solomon packet code is very effective in protecting contiguous nucleotides[17]. In this paper, we use RS coding for error correction where the error correction block size is 240 bits payload with 15 bits of redundancy added, i.e. RS (255, 240, 15).

According to information theory, errors can be categorized into random and erasure errors. This paper uses the forward error correction method to solve the above errors. In this method, the random error is unknown, and the erasure error is known. If *E*_*e*_ is used to denote an erasure error and *E*_*r*_ is used to indicate a random error, RS (255, 240, 15) shall satisfy the following conditions:

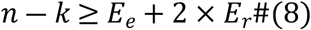

When an insertion or deletion of a base occurs in a DNA sequence, this means that an erasure error has occurred. Since we already know the existence of erasure errors, we must consider the problems posed by substitution errors (random errors) when designing error correction codes. Note that since we did not add external code error correction to the priming and indexing regions, we only analyzed the information bits in the payload region for single-segment reads.

Based on the data in Table 3, we performed reading experiments on three texts, Text 1 (Shakespeare’s Sonnets), Text 2 (DNA Springs Abstracts), and Photograph 1 (Cameraman 256), respectively. For Text 1 (Shakespeare’s sonnets), we synthesized 266,272 nucleotides, from which we selected a fraction of the bases (129 nt × 10 entries, for a total of 1,290 bases) for testing. Substitution errors of 80 bases were randomly added to each DNA sequence, and then automatic recovery experiments were performed. By adding RS (255, 240) error correction codes to the message load portion, we fully recovered it to the original text. For Text 2 (DNA Fountain summary) and photo 1 (Cameraman 256), we carried out the same process as Text 1, and the results are shown in the Table 3.

**Table 3.** Overview of the number of incorporation substitution error nucleotides.

|  | Text 1 | Text 2 | Photo 1 |
| --- | --- | --- | --- |
| Total number of bases in DAN | 266272 | 2255 | 475653 |
| Testing the number of DNA bases | 1290 | 1290 | 1290 |
| Add replacement error | 80 | 2 | 120 |
| Recovery situation | successes | successes | successes |
| decoding ratio | 100% | 100% | 100% |

According to the results in Table 4, among all the generated DNA sequences, we randomly selected 1290 bases (129 nt × 10 entries) for insertion and deletion error simulation experiments. We added three random deletion errors to DNA sequence #8 in Text 1. However, only two deletion errors were corrected in the final decoding, resulting in a decoding rate of 99.84% and an absolute decoding failure. In Text 2 (DNA Fountain Abstracts), the number of errors added during the simulation was low and fully recovered. For photo 1 (Cameraman 256), we performed the same process as for Text 1 and the results are shown in the Table 4.

**Table 4.**
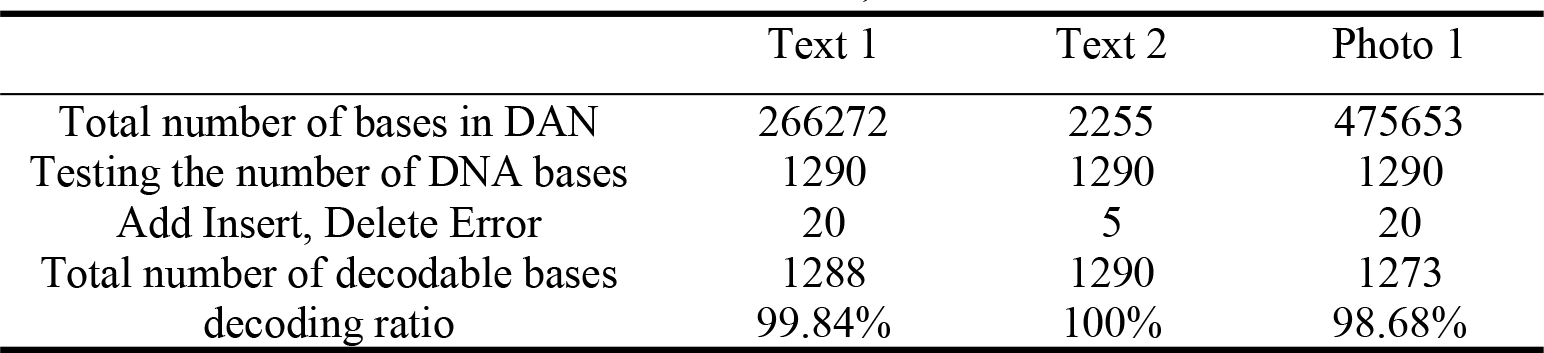
Overview of the number of inserted, deleted error nucleotides added to the.

|  | Text 1 | Text 2 | Photo 1 |
| --- | --- | --- | --- |
| Total number of bases in DAN | 266272 | 2255 | 475653 |
| Testing the number of DNA bases | 1290 | 1290 | 1290 |
| Add Insert, Delete Error | 20 | 5 | 20 |
| Total number of decodable bases | 1288 | 1290 | 1273 |
| decoding ratio | 99.84% | 100% | 98.68% |

## 5 Simulation integration and comparison sequences

### 5.1 Informative DNA primer design

According to the previous coding rules, Shakespeare’s sonnets yield 266,272 bases (1,850 DNA sequences). Short homologs of 129 bases were first synthesized, and then primers were designed using the Oligo 7 program for these short homologs to facilitate subsequent PCR. Considering the DNA sequence design requirements in the previous section and the dissolution temperature requirements in the actual PCR operation, we designed the primer sequences of the short homologs to be 20 bases ± 1 base, Tm = 53.9 °C, and finally created the forward primers and reverse primers as shown in Table 5 (in part).

**Table 5.**
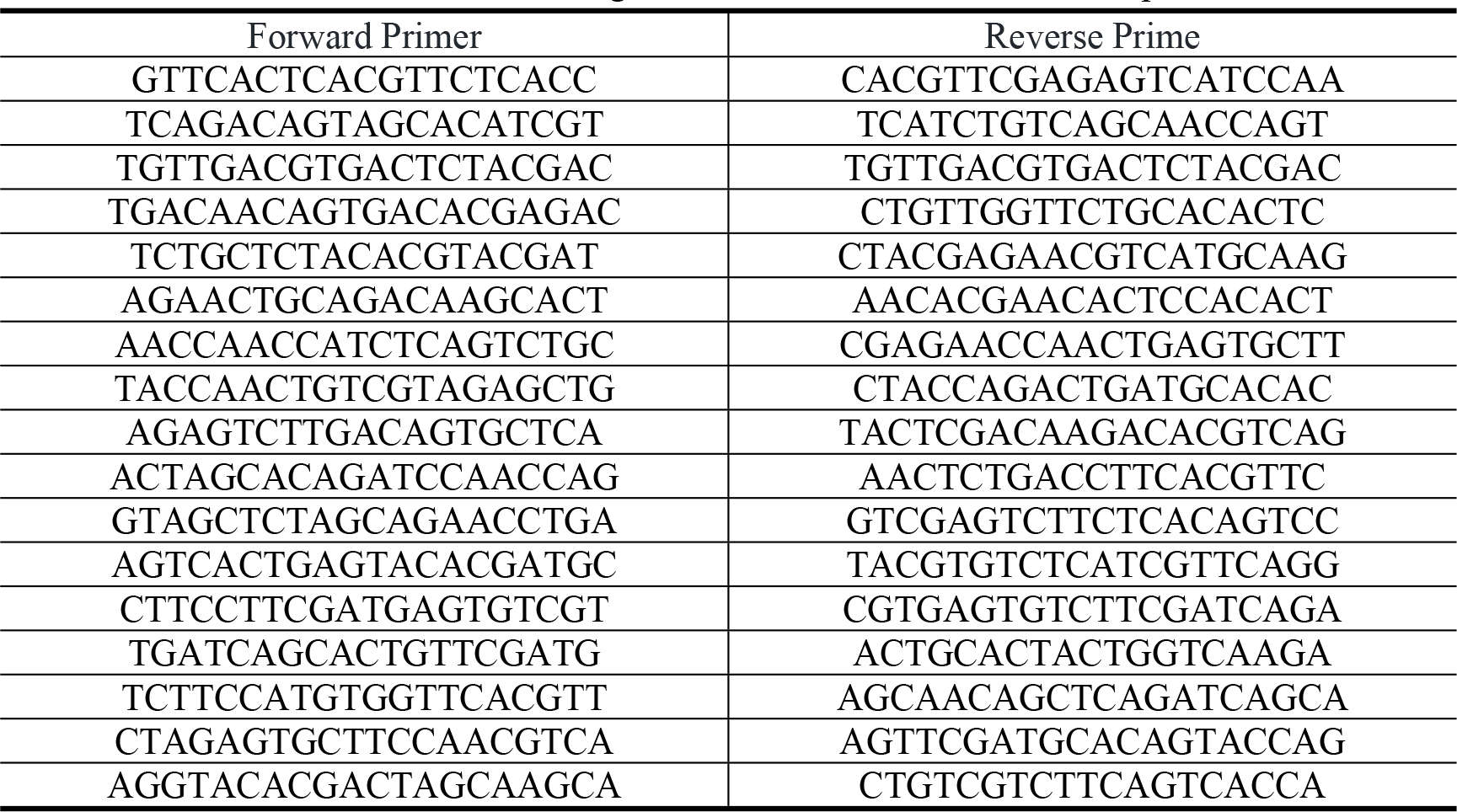
Primer design for forward and reverse DNA sequences.

We performed a GC content test and dissolution temperature Tm statistics for the above primer sequences. The results are shown in Fig. 6(f) and Fig. 7. According to Fig. 6(f), all the designed primer sequences were within the reasonable range of dissolution temperatures (45°C∼65°C). Fig. 7 shows that no primer sequences were found to have GC content exceeding the standard. (The highest percentage of GC content was 56%, and the lowest was 44%)

**Fig. 7.**
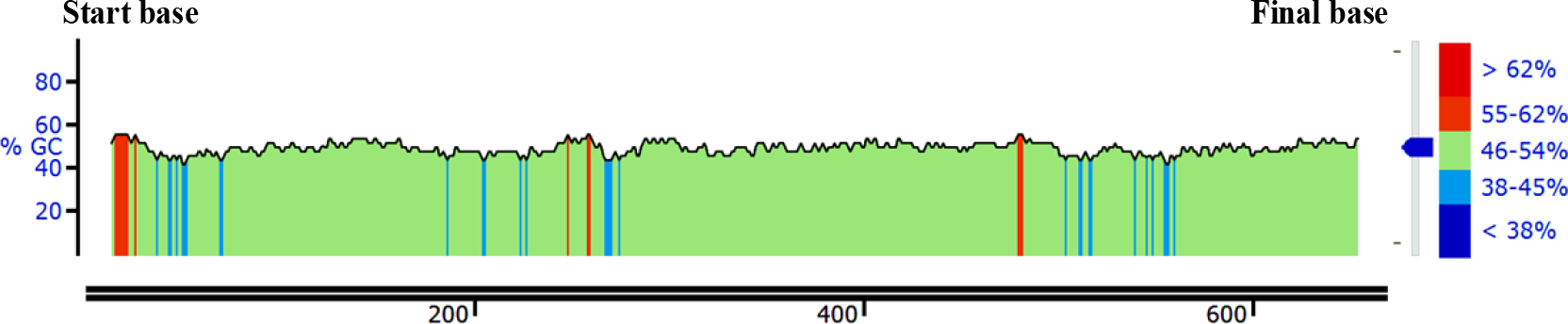
Distribution of GC content in primers

### 5.3 Genomic integration of informational DNA

Recombination of E.coli plasmid vectors in combination with informative DNA, recombinant E.coli plasmid vectors contain resistance genes, integrase genes, and initiation of replication sites along the informative DNA sequence[38]. Then E.coli was chemically transformed. To ensure fidelity when sequencing the informative DNA sequences, we adopted a double-stranded design for the informative DNA sequences, forward sequencing for the informative DNA sequences, and reverse sequencing needed to obtain the sequences consistent with the informative DNA sequences after further transcription. The forward message DNA sequence was digested using EcoR I enzyme, The reverse message DNA sequence was digested using Xba I enzyme, and then the message DNA sequence was integrated into E.coli using T7 DNA ligase, which resulted in the replication of the message DNA along with the value-added of the E.coli plasmid vector[39], as shown in Figure 8.

**Fig. 8.**
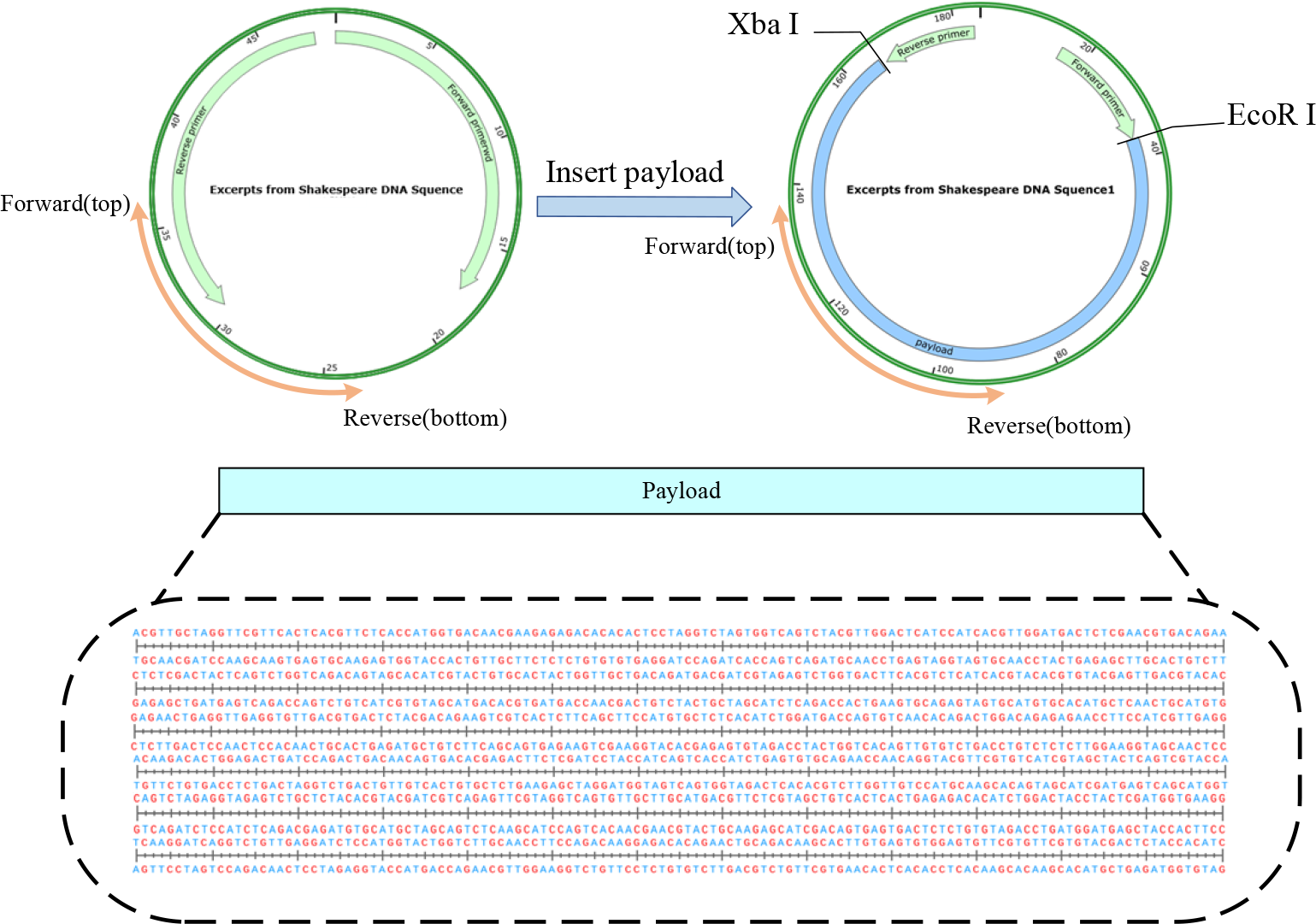
Assembly of informative DNA with E.coli plasmid vector

### 5.4 Simulation of multiple sequence comparison

This study used excerpts from Shakespeare’s sonnets to simulate PCR amplification of DNA sequences. Firstly, according to the previous article, after transcoding the original text into DNA sequences, every 129 bases in the DNA sequences were split into short homologues. The primers for the short homologues were designed according to the Oligo 7 Primer Design Software and finally reorganized into new DNA sequences. We used SnapGene software to simulate the PCR simulation of recombinant DNA sequences by intercepting a short homologue DNA sequence for five PCR amplifications, with the lysis temperature set to Tm = 60 °C. Finally, the PCR-amplified DAN sequences were subjected to multiple sequence comparisons to recover the correct original sequences, and the results are shown in Figure 10.

According to Fig. 9, it can be seen that the encoding and error correction scheme designed in this scheme can completely recover the original information after the DNA columns generated by polymerase chain reaction (PCR) and then decode them after multiple sequence comparisons.

**Fig. 9.**
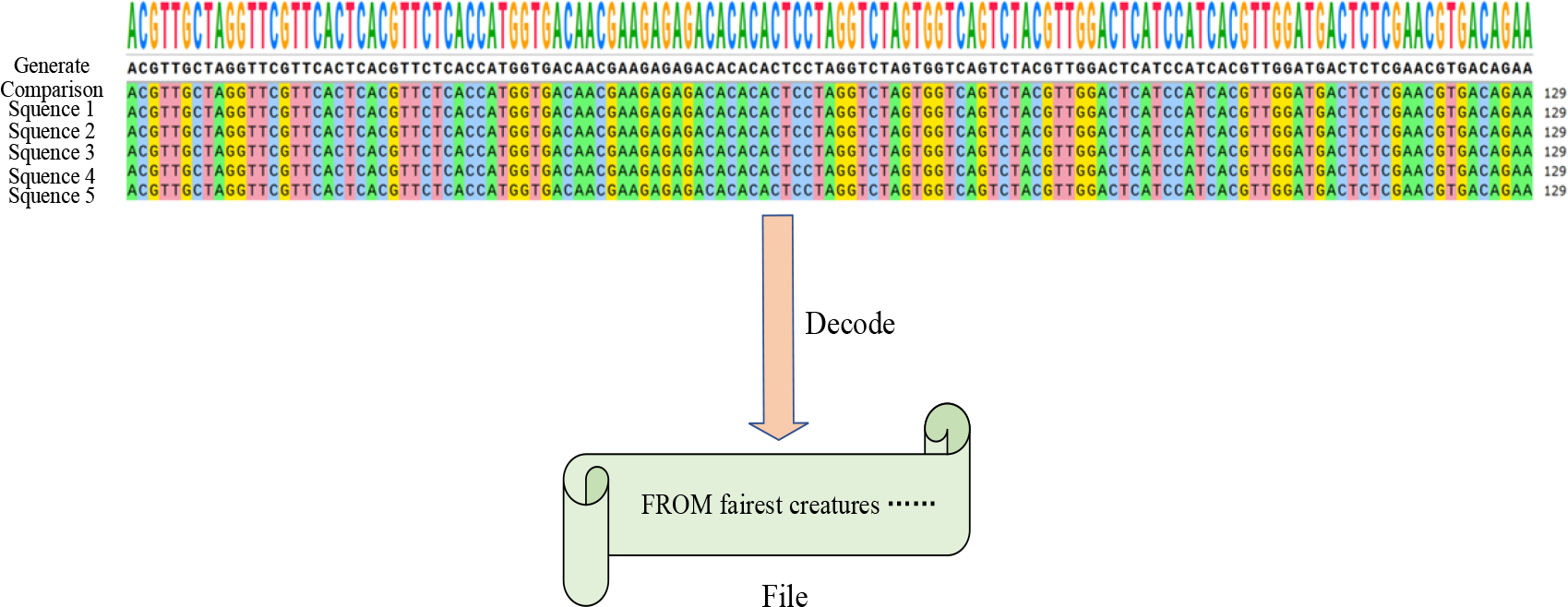
Comparison and decoding of 5 DNA sequences

## 6 Conclusion and Outlook

DNA bases are essential as a potential long-term storage medium for data that needs to be preserved for a long time. DNA molecules can remain stable in dry, cold and dark environments for at least thousands of years, unlike standard large-scale physical media that require expensive data migration fees to adapt to environmental changes. Digital information storage using DNA molecules could theoretically achieve storage densities of 11 EB/mm^3^, and information stored in DNA molecules could be read at any time without risking the obsolescence of the extraction method.

This paper successfully simulates and analyses texts such as Shakespeare’s sonnets for GC content, homopolymer, and error correction performance. The constraints in subsequent biological experiments can be satisfied using the proposed 64-element mapping table. It prevents the increase of errors such as base substitutions, deletions and insertions due to extreme GC content and also contains misreading situations during sequencing due to long homopolymers. This scheme analyzes the error correction and single-segment read performance and provides recovery rate statistics for data reads of text and image types. The shortcoming to be pointed out is that we only added the error correction code in the information loading part of the DNA sequence without the outer code protection of the error correction code for the primer region, which is what we need to improve in the next step. In terms of coding efficiency, the coding storage density of this scheme is 1.49 bit/nt. However, there is still a gap compared with the maximum theoretical storage density of 2 bit/NT per nucleotide. In the follow-up work, we will improve the 64-element coded table and introduce the concept of concatenated bases to enhance the coding density further.

## Notes

### Competing Interest Statement

The authors have declared that no competing interests exist.

## Reference

[1] Y. Hao, Q. Li, C. Fan and F. Wang. “Data Storage Based on DNA,” Small Structures.vol. 2, no. 2, sep, 2020.

[2] X. Zan, X. Yao, P. Xu, Z. Chen, L. Xie, S. Li and W. Liu. “A Hierarchical Error Correction Strategy for Text DNA Storage,” Interdisciplinary Sciences: Computational Life Sciences.vol. 14, no. 1, pp.141–150, Aug, 2022.

[3] Y. Yesiltepe, J. R. Nuñez, S. M. Colby, D. G. Thomas, M. I. Borkum, P. N. Reardon, N. M. Washton, T. O. Metz, J. G. Teeguarden, N. Govind and R. S. Renslow. “An automated framework for NMR chemical shift calculations of small organic molecules,” Journal of Cheminformatics.vol. 10, no. 1, pp.52, Oct, 2018.

[4] M. H. Raza, S. Desai, S. Aravamudhan and R. Zadegan. “An outlook on the current challenges and opportunities in DNA data storage,” Biotechnology Advances.vol. 66, pp.108155, Sep, 2023.

[5] E. L. van Dijk, Y. Jaszczyszyn, D. Naquin and C. Thermes. “The Third Revolution in Sequencing Technology,” Trends in Genetics.vol. 34, no. 9, pp.666–681, Sep, 2018.

[6] L. Organick, S. D. Ang, Y.-J. Chen, R. Lopez, S. Yekhanin, K. Makarychev, M. Z. Racz, G. Kamath, P. Gopalan, B. Nguyen, C. N. Takahashi, S. Newman, H.-Y. Parker, C. Rashtchian, K. Stewart, G. Gupta, R. Carlson, J. Mulligan, D. Carmean, G. Seelig, L. Ceze and K. Strauss. “Random access in large-scale DNA data storage,” Nature Biotechnology.vol. 36, no. 3, pp.242–248, Feb, 2018.

[7] L. Piantanida and W. L. Hughes. “A PCR-free approach to random access in DNA,” Nature Materials.vol. 20, no. 9, pp.1173–1174, Aug, 2021.

[8] F. Sun, Y. Dong, M. Ni, Z. Ping, Y. Sun, Q. Ouyang and L. Qian. “Mobile and Self-Sustained Data Storage in an Extremophile Genomic DNA,” Advanced Science.vol. 10, no. 10, pp.2206201, Feb, 2023.

[9] Y. Erlich and D. Zielinski. “DNA Fountain enables a robust and efficient storage architecture,” Cold Spring Harbor Laboratory.vol. 355.no. 6328, Mar, 2016.

[10] F. Pfeiffer, C. Gröber, M. Blank, K. Händler, M. Beyer, J. L. Schultze and G. Mayer. “Systematic evaluation of error rates and causes in short samples in next-generation sequencing,” Scientific Reports.vol. 8, no. 1, pp.10950, Jul, 2018.

[11] Z. Ping, S. Chen, X. Huang, S. J. Zhu and Y. Shen. “Towards Practical and Robust DNA-based Data Archiving by Codec System Named ‘Yin-Yang’,” Cold Spring Harbor Laboratory.vol.2 no.4, pp. 234–242, Apr, 2022.

[12] J. Koch, S. Gantenbein, K. Masania, W. J. Stark, Y. Erlich and R. N. Grass. “A DNA-of-things storage architecture to create materials with embedded memory,” Nature Biotechnology.vol. 38, no. 1, pp.39–43, Dec, 2020.

[13] L. C. Meiser, P. L. Antkowiak, J. Koch, W. D. Chen and R. N. Grass. “Reading and writing digital data in DNA,” Nature Protocols.vol. 15, no. 1, pp. 86–101, Nov, 2019.

[14] M. Welzel, P. M. Schwarz, H. F. Löchel, T. Kabdullayeva, S. Clemens, A. Becker, B. Freisleben and D. Heider. “DNA-Aeon provides flexible arithmetic coding for constraint adherence and error correction in DNA storage,” Nature Communications.vol. 14, no. 1, pp.628, Feb, 2023.

[15] G. M. Church, Y. Gao and S. Kosuri. “Next-Generation Digital Information Storage in DNA,” Science.vol. 337, no. 6102, pp.1628, Aug, 2012.

[16] R. N. Grass, R. Heckel, M. Puddu, D. Paunescu and W. J. Stark. “Robust Chemical Preservation of Digital Information on DNA in Silica with Error-Correcting Codes,” Angewandte Chemie.vol.54. no.8, Feb, 2015.

[17] M. Blawat, K. Gaedke, I. Huetter, X. M. Chen, B. Turczyk, S. Inverso, B. Pruitt and G. Church. “Forward Error Correction for DNA Data Storage,” Procedia Computer Science.vol. 80, pp.1011–1022, 2016.

[18] Y. Erlich and D. Zielinski. “dna storage dna fountain enables a robust and efficient storage architecture,’’vol.355. no.6328. pp. 950–954, Mar, 2018.

[19] L. Feng, C. H. Foh, J. Cai and L. T. Chia. “LT codes decoding: design and analysis,” 2009 IEEE International Symposium on Information Theory.vol. no. 2009.

[20] K. Zhang, J. Jiao, S. Wu and Q. Zhang. “Short Analog Fountain Code With Quasi-Gray Constellation Mapping Modulation Towards uRLLC,” IEEE Transactions on Signal Processing.vol. 70, pp.4077–4092, 2022.

[21] L. Organick, Y.-J. Chen, S. Dumas Ang, R. Lopez, X. Liu, K. Strauss and L. Ceze. “Author Correction: Probing the physical limits of reliable DNA data retrieval,” Nature Communications.vol. 11, no. 1, pp.1080, 2020.

[22] A. S. Tanenbaum and H. Bos. “Modern operating systems, Fourth Edition,’’vol. no. 2015.

[23] ,“Towards practical and robust DNA-based data archiving using the yin–yang codec system,” Nature Computational Science.vol.2. no.4.2022

[24] M. Hao, H. Qiao, Y. Gao, Z. Wang, X. Qiao, X. Chen and H. Qi. “A mixed culture of bacterial cells enables an economic DNA storage on a large scale,” Communications Biology.vol.3. no.1.pp. 1–9. 2020

[25] W. H. Press, J. A. Hawkins, S. K. Jones, J. M. Schaub and I. J. Finkelstein. “HEDGES error-correcting code for DNA storage corrects indels and allows sequence constraints,” Proceedings of the National Academy of Sciences.vol. 117, no. 31, pp.18489–18496, 2020.

[26] S. J. Park, Y. Lee and J. S. No. “Iterative coding scheme satisfying GC balance and run-length constraints for DNA storage with robustness to error propagation,” Journal of Communications and Networks.vol. 24, no. 3, pp.283–291, 2022.

[27] C. Ezekannagha, M. Welzel, D. Heider and G. Hattab. “DNAsmart: Multiple attribute ranking tool for DNA data storage systems,” Computational and Structural Biotechnology Journal.vol. 21, pp.1448–1460, 2023.

[28] A. T. Duckworth, K. Bilotti, V. Potapov and G. J. S. Lohman. “Profiling DNA Ligase Substrate Specificity with a Pacific Biosciences Single-Molecule Real-Time Sequencing Assay,” Current Protocols.vol. 3, no. 3, pp.e690, 2023.

[29] X. Lu, Y. Huang, S. Yan, W. Yang and P. Haskell-Dowland. “Energy-Efficient Covert Wireless Communication Through Probabilistic Jamming,” IEEE Wireless Communications Letters.vol. 12, no. 5, pp.932–936, 2023.

[30] R. Heckel, G. Mikutis and R. N. Grass. “A Characterization of the DNA Data Storage Channel,” Scientific Reports.vol.9. no.1. 2019.

[31] Y. Dong, F. Sun, Z. Ping, Q. Ouyang and L. Qian. “DNA storage: research landscape and future prospects,’’vol. 7, no. 6, pp.1092–1107, 2020.

[32] J. Koch, S. Gantenbein, K. Masania, W. J. Stark and R. Grass. “SI Video for DNA-of-things storage architecture to create materials with embedded memory,” Nature Biotechnology.vol. 38, no. 1, 2020.

[33] M. F. Laursen, M. D. Dalgaard and M. I. Bahl. “Genomic GC-Content Affects the Accuracy of 16S rRNA Gene Sequencing Based Microbial Profiling due to PCR Bias,” Frontiers in Microbiology.vol. 8, pp.1934, 2017.

[34] S. Yazdi, R. Gabrys and O. Milenkovic. “Portable and Error-Free DNA-Based Data Storage,” Cold Spring Harbor Laboratory.vol.7. no. 1, 2017.

[35] N. Goldman, P. Bertone, S. Chen, C. Dessimoz, E. M. Leproust, B. Sipos and E. Birney. “Towards practical, high-capacity, low-maintenance information storage in synthesized DNA,” Nature.vol. 494, no. 7435, pp.77–80, 2013.

[36] J. Bornholt, R. Lopez, D. M. Carmean, L. Ceze and K. Strauss. “A DNA-Based Archival Storage System,” IEEE Micro.vol.50, no. 2, pp.637–649, 2016.

[37] M. Kolmogorov, J. Yuan, Y. Lin and P. A. Pevzner. “Assembly of long, error-prone reads using repeat graphs,” Nature Biotechnology.vol. 37, no. 5, pp.540–546, 2019.

[38] S. Zucca, L. Pasotti, N. Politi, M. G. Cusella De Angelis and P. Magni. “A standard vector for the chromosomal integration and characterization of BioBrick™ parts in Escherichia coli,” Journal of Biological Engineering.vol. 7, no. 1, pp.12, 2013.

[39] E. Falgenhauer, S. von Schönberg, C. Meng, A. Mückl, K. Vogele, Q. Emslander, C. Ludwig and F. C. Simmel. “Evaluation of an E. coli Cell Extract Prepared by Lysozyme-Assisted Sonication via Gene Expression, Phage Assembly and Proteomics,” ChemBioChem.vol. 22, no. 18, pp.2805–2813, 2021.

